# Matrix elasticity gradients guide neuronal polarity by controlling microtubule network mobility

**DOI:** 10.1101/2021.07.22.453336

**Authors:** Mithila Burute, Klara I. Jansen, Marko Mihajlovic, Tina Vermonden, Lukas C. Kapitein

## Abstract

Neuronal polarization and axon specification depend on extracellular cues, intracellular signaling, cytoskeletal rearrangements and polarized transport, but the interplay between these processes has remained unresolved. The polarized transport of kinesin-1 into a specific neurite is an early marker for axon identity, but the mechanisms that govern neurite selection and polarized transport are unknown. We show that extracellular elasticity gradients control polarized transport and axon specification, mediated by Rho-GTPases whose local activation is necessary and sufficient for polarized transport. Selective Kinesin-1 accumulation furthermore depends on differences in microtubule network mobility between neurites and local control over this mobility is necessary and sufficient for proper polarization, as shown using optogenetic anchoring of microtubules. Together, these results explain how mechanical cues can instruct polarized transport and axon specification.

## RESULTS AND DISCUSSION

The proper functioning of neurons depends on their polarized organization into axons and dendrites. Neuronal polarization and subsequent axon specification require a delicate coordination between extracellular cues, intracellular signaling, cytoskeletal rearrangements and selective transport into specific neurites, but the precise interplay between these processes is not known (*1*–*3*). Recent work revealed the importance of extracellular mechanical cues in guiding axonal growth, but whether neuron polarization and axon specification can be controlled by the mechanical properties of the extracellular matrix is unknown (*4*).

Importantly, dissociated neurons that lack spatially-defined extracellular cues also properly polarize (*5*). Several hours after plating these neurons (stage 1) they grow multiple neurites (stage 2), of which one becomes the axon (stage 3), whereas the others later develop into dendrites (stage 4). In these neurons, the selective accumulation of the microtubule-based motor protein Kinesin-1 is an early marker for the future axon (*6*–*8*). Remarkably, before accumulating in the future axon, Kinesin-1 first transiently enriches in a small and alternating subset of neurites during stage 2, reflecting some form of transient polarization (Fig. S1 and Movie S1) (*6, 8, 9*). Various models have been proposed for this polarized transport of Kinesin-1, but none of these explain how Kinesin-1 cycles between neurites and how this cycling can be modulated to more precisely define the future axon (*3, 7, 10*).

We set out to dissect the interplay between mechanical cues, intracellular signaling and microtubule rearrangements in guiding polarized transport and axon formation. To explore whether the mechanical properties of the extracellular matrix (ECM) can guide axon specification, we monitored axon formation in dissociated embryonic hippocampal neurons grown on Laminin-coated gels with a stiffness varying from 0.4–6 kPa (Fig. S2). We used a recently introduced method to generate gradient gels in which the density of embedded fluorescent beads reports the local stiffness and can be used to map the stiffness landscape surrounding each neuron (Fig. S2) (*11*). Dissociated Rat hippocampal neurons were plated on these gels directly upon dissection and were fixed and stained for the axonal protein Tau after 2 days. This revealed that ∼82% of axons in stage 3 neurons were directed towards the softer side of the matrix (Fig. 1, A toC). Tracing the stiffness profile along each axon yielded an average decrease in stiffness of 2 Pa/μm (Fig. 1,D and E, Fig. S3). We furthermore tested whether stiffness gradients could also bias the polarized transport of Kinesin-1 in morphologically unpolarized stage 2 neurons. Indeed, in ∼75% of cells the neurite positive for Kinesin-1 was directed towards the softer side of the gel and the average stiffness gradient along these neurites was 3.7 Pa/μm (Fig. 1 C toE). These results demonstrate that polarized transport, axon specification and axon growth can be guided by stiffness gradients.

**Figure 1.**
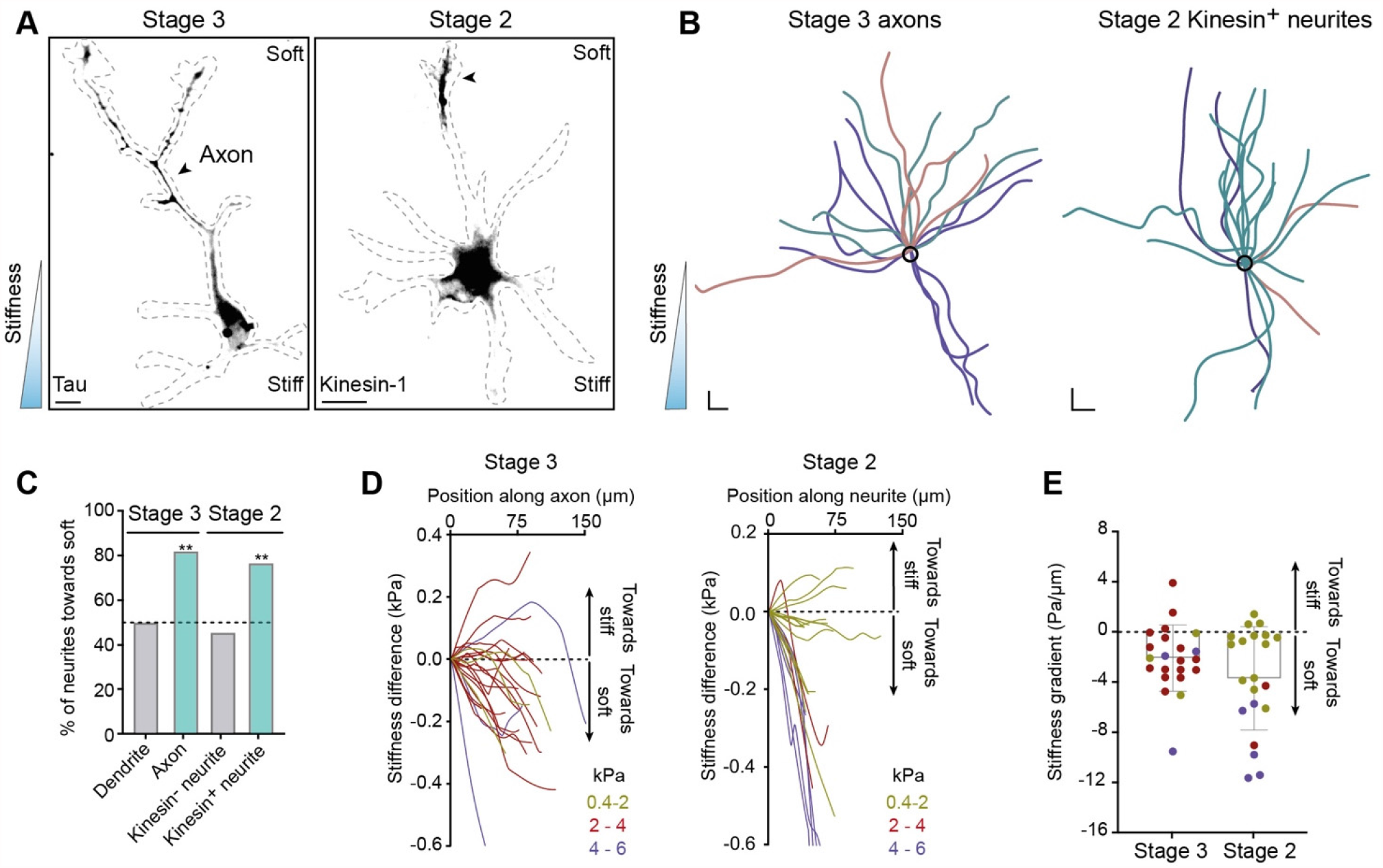
Elasticity gradients control the polarized accumulation of Kinesin-1 and axon formation. (A) Rat embryonic hippocampal neurons on gradient elasticity substrates ranging 0.1-3 kPa. Left: stage 3 neuron labelled for the axon marker Tau. Right: stage 2 neuron expressing Kinesin-1 (K353-GCN4-GFP). Arrowheads mark a neurite with Kinesin-1 accumulation or a Tau-positive axon, both growing towards the softer matrix. Dashed lines indicate neuron morphology. Scale bar, 10 µm. (B) Morphology of stage 3 Tau positive axons (left, n=20) and stage 2 neurites with Kinesin-1 accumulation (right, n=20), aligned with respect to the stiffness gradient. Black circle represents position of cell bodies. Colors represent traces from independent experiments. Scale bar 20 µm (*x,y*). (C) Percentage of stage 3 axons and dendrites (n=22 and 46, N=3 independent experiments) and stage 2 neurites with and without Kinesin-1 accumulation (n=22 and 62 neurites N=3 independent experiments) directed toward the soft matrix on stiffness gradient. ** p≤0.01 (Chi-square test). (D) Stiffness difference relative to the cell body, along the length of stage 3 axons (n=22) and stage 2 neurites with Kinesin-1 (n=21). Colors represent different rigidity ranges for the matrix near the cell body. (E) Overall stiffness gradient for the traces shown in D. Error bars represent S.D.

Mechanical cues from the ECM are often translated into intracellular effects through Rho-GTPase-based signaling, and both Rac1 and Cdc42 have been implicated in neuronal polarity and axon formation (*12*–*15*). To test how these Rho-GTPases affect the mechanically-guided polarized transport of Kinesin-1, we inhibited Rac1 activity in stage 2 neurons grown on stiffness gradient gels and found that the biased accumulation of Kinesin-1 towards the softer side of the gel was lost upon inhibition (Fig. 2, A and B). In stage 2 neurons that were grown on glass, inhibition of either Rac1 or Cdc42 also prevented the selective accumulation of Kinesin-1, which instead enriched at the tip of all neurites (Fig. 2, C, D and Movie S2). Live-imaging confirmed that Rac1 inhibition readily stopped Kinesin-1 cycling between neurites, followed by accumulation in multiple tips, suggesting that local enrichment of Rac1 activity is necessary for selective Kinesin-1 transport (Fig. 2, E and F). We therefore performed local photoactivation of Rac1 and found that this was sufficient to guide Kinesin-1 transport into a single neurite, demonstrating the importance of local activities of axon-promoting Rho-GTPases for polarized transport in stage 2 neurons (Fig. 2, G to I, Fig. S4 and Movie S3).

**Figure 2.**
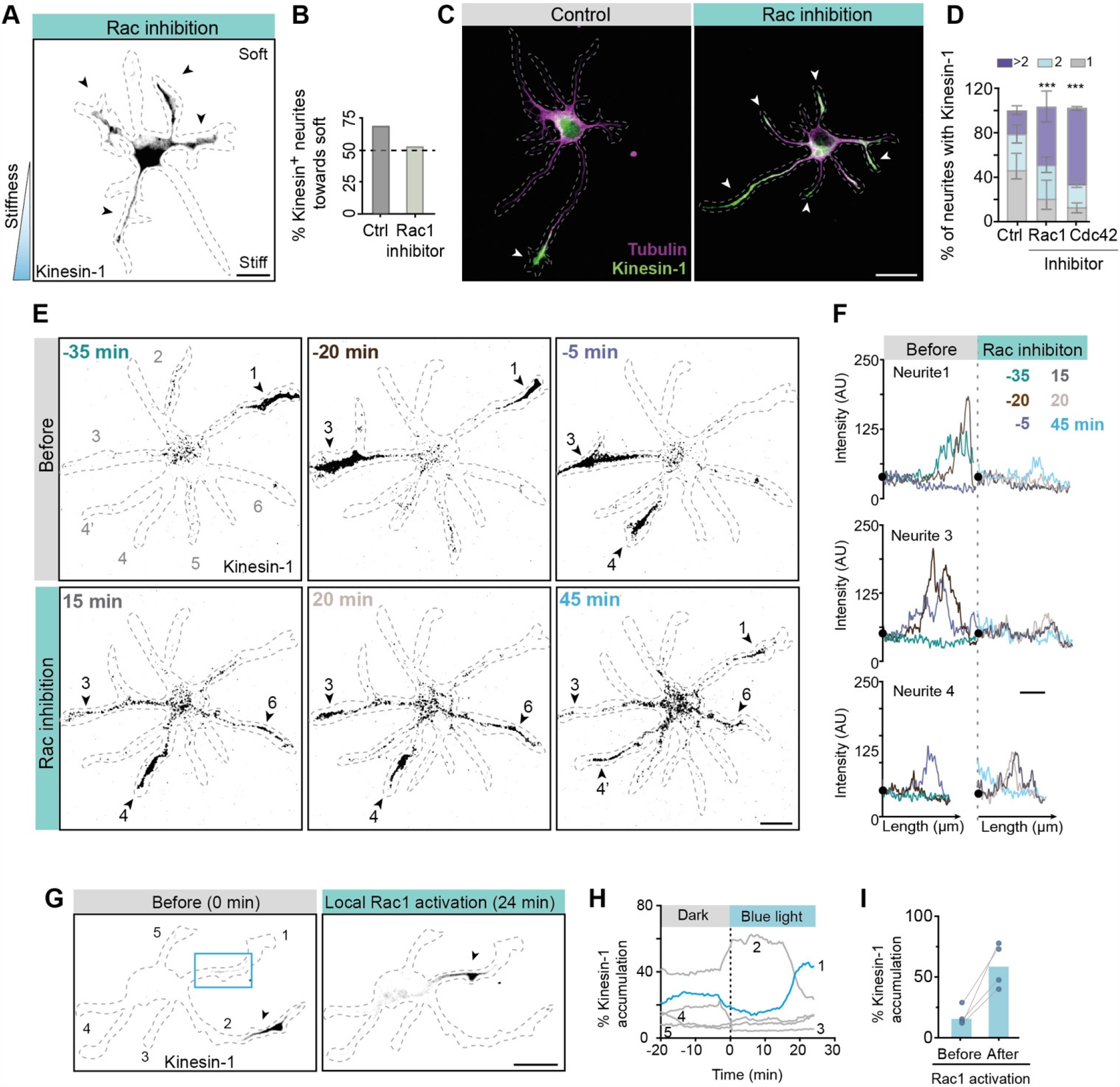
Axon specifying Rho GTPases control selective Kinesin-1 accumulation. (A) Immunofluorescence of stage 2 neuron expressing Kinesin-1, grown on a stiffness gradient matrix and treated with a Rac inhibitor. (B) Percentage of stage 2 neurites with Kinesin-1 enrichment that are positioned towards softer matrix in control conditions (n=21) or upon Rac1 inhibition (n=54) (N=2 independent experiments). (C) Immunofluorescence of stage 2 neurons expressing Kinesin-1. Kinesin 1 mostly accumulated in a single neurite in control conditions (left), but in multiple neurites upon inhibition of Rac1 (right). (A-C) Arrowheads indicate multiple neurites with Kinesin-1 accumulation. (D) Percentage of stage 2 neurons with Kinesin-1 accumulation in 1, 2 or >2 neurites. Control (n=148, Rac inhibitor (n=78), Cdc42 inhibitor (n=77) (N=3 independent experiments). Error bars represent S.D. *** p≤0.001 (Chi-square test). (E) Selected frames from a time-lapse movie (<u>Movie S2</u>) of a stage 2 neuron expressing Kinesin-1. Top three frames show change of selective accumulation of Kinesin-1 from neurite 1 (−35’) to neurite 3 (−20’) and to neurite 3 and 4 (−5’). The (transient) selective accumulation of Kinesin-1 is lost upon inhibition of Rac1 resulting in its sustained accumulation in neurites 1, 3, 4 and 6. (F) Intensity profiles along the length of neurites 1, 3 and 4 for 6 frames shown in (A). Colors are assigned to different time frames. Black dot represents the position of cell body. A.U., arbitrary units. (E)-(F) The Rac inhibitor was added at time point 0’ which is indicated by the dotted line in (F). (G) Kinesin-1 is selectively accumulated in the neurite 2 of stage 2 neuron upon local photoactivation of Rac1. Blue box indicates blue light-photoactivated region. Arrowheads indicate that the Kinesin-1 accumulation changes from neurite 2 to neurite 1 upon Rac1 activation. (H) Percentage of Kinesin-1 accumulation within a neurite is measured for different frames during Rac1 photoactivation. Blue trace indicates the neurite 1 shown in (G) in which Rac1 was activated and gray traces indicated non-activated neurites as numbered in (G). (I) Percentage of Kinesin-1 accumulation in a neurite before and after Rac1 photoactivation in stage 2 neurons (n=4, N=2 independent experiments). Blue bars indicated the median value for measurements. (A)-(G) Neuron morphology is marked by the dashed lines. Scale bar, 20 µm.

Rho-GTPases are known regulators of the cytoskeleton (*16*). To understand how they control the polarized transport of Kinesin-1, we searched for changes in the microtubule cytoskeleton that were associated with changes in the polarized transport of Kinesin-1. Live-cell imaging of the microtubule-associated protein TRIM46, an early axon marker, revealed that its selective localization at the base of specific neurites was strongly correlated with Kinesin-1 transport into that neurite (Fig. 3, A to C, Fig. S5 and Movie S4) (*17*). In addition, more rapid imaging revealed that the various patches of TRIM46 underwent collective anterograde or retrograde motility at 0.04–0.3 μm/s, corresponding with Kinesin-1 entry into or exit from these neurites, respectively (Fig. 3, D and E, and Movie S5). We therefore used local photoactivation to compare the mobility of the microtubule network itself between different neurites (*18*) (Fig. 3, F, G and Movie S6). This revealed that the microtubule network was mostly moving retrogradely in stage 2 neurites, but with occasional reversals in a subset of neurites (Fig. 3, G). In stage 3 neurons, the axon displayed anterograde motility, whereas retrograde motility was observed in the other neurites. During subsequent maturation, the overall mobility of microtubules and TRIM46 dampened within axons (Fig. 3, G and H, Fig. S6, Fig. S7 and Movie S7). To directly correlate microtubule network mobility and Kinesin-1 accumulation, we next combined local tubulin photoactivation with Kinesin-1 imaging. This revealed that exit of Kinesin-1 correlated with retrograde microtubule mobility, whereas Kinesin-1 was retained in the neurite with anterograde microtubule mobility (Fig. 3, I and Movie S8).

**Figure 3.**
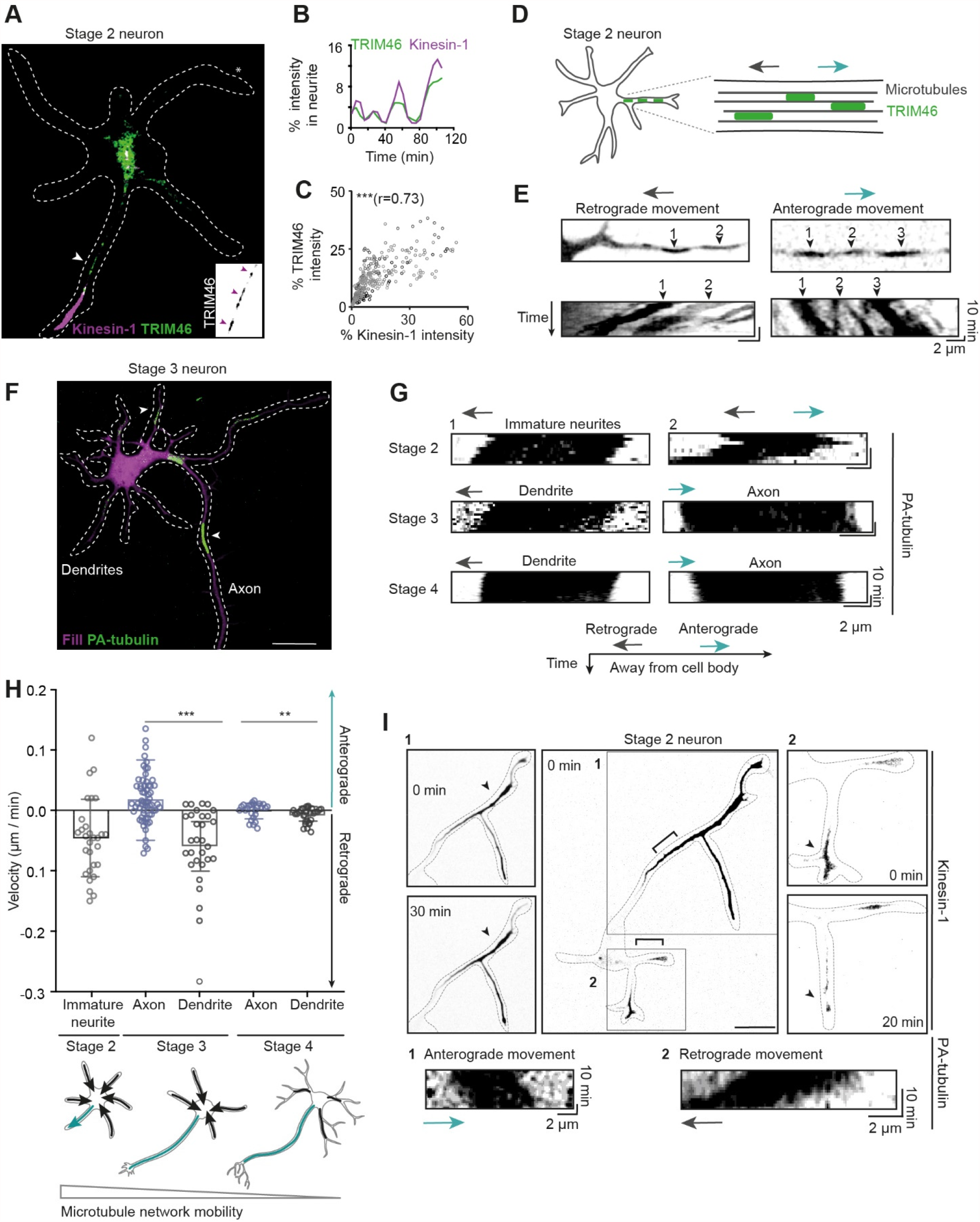
Changes in polarized Kinesin-1 transport correlated with changes in microtubule network mobility. (A) Appearance of TRIM46 in a neurite (white arrowhead) is associated with selective transport of Kinesin-1. Inverted grey inset shows TRIM46 patches (magenta arrowheads) in the neurite. (B) Change in percentage of intensity of TRIM46 and Kinesin-1 in the neurite marked with * in (A). (C) Correlation between TRIM46 and Kinesin-1 enrichment in neurites (n=322 *xy* pairs, 20 neurites from 4 neurons). Grey shades represent data from different neurons. *** p≤0.001. r is Peasron correlation coefficient (Pearson correlation test). (D-E). Directional movement of TRIM46 patches towards (anterograde) and away (anterograde) from cell body. Top images show start position and kymographs show movement of TRIM46 patches. (F) Stage 3 neuron expressing mRFP-fill (magenta) and Photoactivable-tubulin (PA-tubulin). Arrowhead indicate PA-tubulin regions (green) in axon and dendrite. (G) Kymographs representing movement of PA-tubulin regions at stage 2-4 neurons. (H) Top: velocities of anterograde (positive values) and retrograde (negative values) microtubule movement for stage 2 neurites (n=29 regions), stage 3 dendrites (n=30 regions) and axons (n=62 regions), stage 4 axons (n=25 regions) and dendrites (n=28 regions). ***P≤0.001, **≤0.01 (Mann Whitney test). Bottom: summary of microtubule network mobility at different stages. (I) Stage 2 neuron showing correlated changes in Kinesin-1 accumulation and PA-tubulin movement. Neurite 1 has an anterograde MT network mobility and retains Kinesin-1 during 30 min (two left images). Neurite 2 has a retrograde network mobility and loses Kinesin-1 within 20 min (two right images).

Together, these results indicate that stiffness gradients control selective Kinesin-1 transport by local control over microtubule mobility, suggesting that such gradients cannot bias Kinesin-1 accumulation if the microtubule network is globally immobilized. To test this, we used chemically-induced dimerization to globally anchor microtubules to the cell membrane (Fig. 4, A). In our approach, an FKBP domain was targeted to the plasma membrane (FKBP-CaaX), where addition of the small molecule Rapalog would trigger interaction with its binding partner FRB fused to a Kinesin mutant known to tightly interact with microtubules (*19*). We validated this approach in non-neuronal cells, where individual microtubules were found to robustly enrich near the plasma membrane within 40 minutes upon Rapalog addition and displayed reduced movements (Fig. S8 and Movie S9). When used in stage 2 neurons plated on substrates with stiffness gradients, selective Kinesin-1 transport was lost upon Rapalog treatment and as a consequence, Kinesin-1 accumulation was no longer biased towards the softer matrix (Fig. 4, B and C).

**Figure 4.**
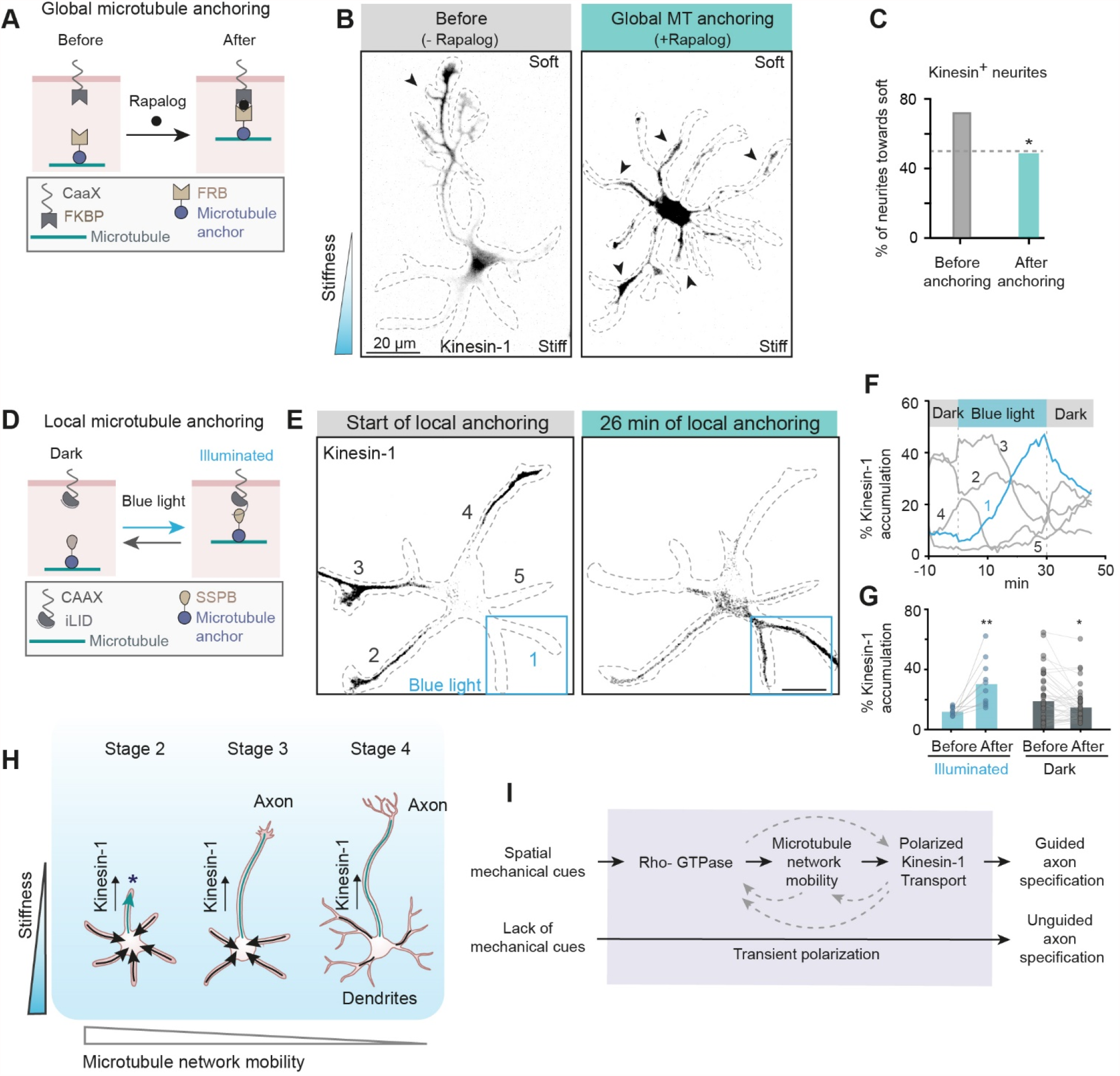
Controlling polarized transport by modulating microtubule network mobility. (A) Rapalog-induced global microtubule anchoring assay. FKBP-FRB dimerization is induced by addition of Rapalog, resulting in global anchoring of microtubules to the cell membrane. (B) Immunofluorescence of stage 2 neuron growing on stiffness gradient. Without Rapalog addition, neurites with Kinesin-1 accumulation are oriented towards softer matrix (left image), whereas Kinesin-1 is accumulated in multiple neurites after global microtubule anchoring (right image). (C) Percentage of stage 2 neurites with Kinesin-1 accumulation towards softer matrix, before (n=43, and after Rapalog addition (n=63) (N=2 independent experiments). *p≤0.05 (Chi-square test). (D) Light-induced local microtubule anchoring assay. Blue light induces binding between iLID-CaaX and SspB-microtubule anchor, resulting in local anchoring of microtubules to the cell membrane. (E) Stage 2 neuron before and after local blue-light illumination (in blue rectangle). Kinesin-1 is relocated from neurite 2, 3, 4 at 0 min to neurite 1 at 26 min. (F) Time traces of Kinesin-1 accumulation in multiple neurites of the neuron shown in D. The blue trace represents the neurite illuminated by blue light. (G) Quantification of Kinesin-1 accumulation before and after blue-light activation in neurites activated for 40 min (blue dots) (n=10 pairs) and non-illuminated neurites in the same neurons (grey dots) (n=42 pairs) (N=3 independent experiments). Rectangles represent median values. **P≤0.01, *P≤0.05 (two tailed Wilcoxon matched-pairs test). (H) Graphical summary depicting interplay between substrate elasticity, microtubule mobility and Kinesin-1 accumulation during development. (I) Model: Mechanical cues from the ECM activate local Rho-GTPase signaling, resulting in differential MT network mobility between neurites to guide sustained Kinesin-1 polarized transport. Solid arrows depict our experimentally established dependencies, dashed lines depict potential additional dependencies and feedback mechanisms. In the absence of spatial mechanical cues from ECM, selective transport to Kinesin-1 to a neurite is transient. Scale bars, 20 µm.

Because global anchoring of microtubules triggered Kinesin-1 accumulation in all neurites, we next tested whether local immobilization of microtubules by local anchoring was sufficient to control the selective transport of Kinesin-1. To this end, we exchanged the modules for chemically-induced dimerization for modules that enable light-induced dimerization, i.e. iLID and SspB (*20*) (Fig. 4, D). Within 30 minutes after light-induced local anchoring of the microtubule network, Kinesin-1 transport was robustly guided into the selected stage 2 neurite along with enrichment of TRIM46 (Fig. 4, E and F, Fig. S9, Movie S10 and Movie S11). In contrast, when the microtubule anchor was expressed without SspB, local illumination did not alter the polarized transport of Kinesin-1 (Fig. S10 and Movie S12). Thus, local control over microtubule mobility is sufficient to control polarized transport.

In this work, we demonstrated that extracellular elasticity gradients control the polarized transport of Kinesin-1 and subsequent axon specification, mediated by the activity of Rac1 and Cdc42 GTPases (Fig. 4, H and I). Recent work has shown that RhoA activity increases with matrix stiffness, resulting in reduced growth of neurites (*21*). In the presence of stiffness gradients, neurites interacting with the softer matrix would have lower RhoA activity, which could promote Rac1/Cdc42 activity on the softer side through mutual inhibition feedbacks and thereby bias polarized transport and axon formation (*22, 23*).

We furthermore found that the polarized transport of Kinesin-1 depends on the dynamics of the microtubule network within neurites. Retrograde movement of the microtubule network results in Kinesin-1 exit and global microtubule immobilization is sufficient to prevent such exit, whereas neurite-specific microtubule immobilization using optogenetics can be used to control polarized transport and establish accumulation in a neurite of choice (Fig. 4). What causes the rapid redistribution of Kinesin-1 upon local microtubule anchoring or local activation of Rac1? The observed changes in microtubule network mobility (∼0.1 µm/min) are too subtle to directly explain the complete loss of Kinesin-1 from a neurite, because Kinesin-1 itself moves >500 times faster and would easily outrun such retrograde movement (*24*). We hypothesize that Kinesin-1 activity is not just dictated by the organization of the microtubule network, but also actively controlled by Rho-GTPases. In addition, the interplay between Rho-GTPases, microtubules and motors may be further strengthened by various forms of feedback, including the feedback from Kinesin-1 transport to Rho-GTPase signaling (*25*), from Kinesin-1 to microtubule conformation and organization (*10, 26, 27*), and from microtubule network mobility to local Rho-GTPase signaling (*28, 29*) (Fig. 4, I).

Our work reveals the cellular cascade by which extracellular mechanical cues specify the polarity axis of neurons. During brain development, the mechanical properties of the ECM are dynamic and can rapidly change (*30, 31*). Here, we report changes in microtubule network mobility and polarized transport within tens of minutes, suggesting that the microtubule cytoskeleton can rapidly rearrange in response to dynamic mechanical signals from the ECM in order to control polarized transport and axon specification. This newly identified interplay between the ECM, the microtubule cytoskeleton and polarized transport that may be crucial for correct axon orientation and outgrowth during brain development.

## ACKNOWLEDGEMENTS

We thank Anna Akhmanova for constructive feedback and Eugene Katrukha for helpful discussions about image analysis. We thank Max Adrian for generating the Lifeact-mCherry construct. This work was supported by EMBO long-term fellowship (EMBO ALTF 407-2017), Netherlands Organization for Scientific Research (NWO-Graduate program project 022.006.001) and the European Research Council (ERC Consolidator Grant 819219).

## AUTHOR CONTRIBUTIONS

M.B and L.K conceptualized the study. M.B. performed most of the experiments and analyses. K.J. generated DNA constructs, performed experiments for Fig. S8 and assisted with other experiments. M.B and L.K wrote the manuscript. M.M and T.V performed rheology measurements. L.K. supervised the study.

## SUPPLEMENTAL INFORMATION

### MATERIAL AND METHODS

#### Table of chemicals

| Chemicals | Source | Reference |
| --- | --- | --- |
| 16% paraformaldehyde EM grade | Electron microscopy Sciences | Cat #15710 |
| 2% Bis-acrylamide solution | Fischer Scientific | Cat #BP1404 |
| 3-(trimethoxysilyl)propyl methacrylate | Sigma-Aldrich | Cat #M6514 |
| 40% Acrylamide solution | Sigma-Aldrich | Cat #A4058 |
| 8% Glutaraldehyde solution | Sigma-Aldrich | Cat #G7526 |
| Acetic acid (J. T. Baker) | Fisher scientific | Cat # 15578364 |
| Alexa Fluor Plus 405 Phalloidin | Thermo Fisher Scientific | Cat #A30104 |
| Ammonium persulfate | Sigma | Cat #215589 |
| B27 supplement | Gibco | Cat #17504001 |
| Bovine serum albumin (BSA) | Carl Roth | Cat #8076.3 |
| DMSO, anhydrous | Thermo Fisher Scientific | Cat #D12345 |
| EHT1864 (Rac inhibitor) | TOCRIS | Cat #872 |
| Ethanol | Fisher Scientific | Cat #15568474 |
| FluoSpheres Carboxylate-Modified Microspheres, 0.2 um, dark red fluorescent (660/680), 2% solids | Thermo Fischer Scientific | Cat #F8807 |
| FuGENE 6 | Promega | Cat #E2692 |
| L-glutamic acid | Sigma-Aldrich | Cat #G1251 |
| L-glutamine | Gibco | Cat #25030-081 |
| Hydrazine hydrate | Sigma-Aldrich/Merck | Cat #225819 |
| Laminin | Roche | Cat #1124321700 |
| Leibovitz's L-15 medium | Thermo Fischer Scientific | Cat #11415064 |
| Lipofectamine 2000 | Thermo Fischer Scientific | Cat #11668019 |
| ML141(Cdc42 inhibitor) | TOCRIS | Cat #4266 |
| Neurobasal medium | Gibco | Cat #21103049 |
| PBS | Lonza | Cat #BE17-517Q |
| Penicillin/Streptomycin | Gibco | Cat #15140122 |
| Poly-L-Lysine hydrobromide | Sigma-Aldrich | Cat #P2636 |
| Prolong Diamond antifade mountant | Thermo Fischer Scientific | Cat #P3670 |
| Rapalog | Takara | Cat #635057 |
| TEMED | Bio-Rad | Cat #610801 |

#### Table of antibodies

| **Antibody** | **Source** | **Reference** |
| --- | --- | --- |
| Goat anti-mouse Alexa Fluor 488 | Life Technologies | Cat #A11029 |
| Goat anti-mouse Alexa Fluor 568 | Life Technologies | Cat # A11031 |
| Goat anti-Rabbit Alexa Fluor 488 | Life Technologies | Cat # A11034 |
| Goat anti-rat Alexa Fluor 594 | Life Technologies | Cat #A11007 |
| Mouse anti Tau | Merck | Cat #MAB3420 |
| Mouse anti-tubulin acetylated | Sigma-Aldrich | Cat #T7451 |
| Phalloidin Alexa Fluor 488 | Thermo Fisher Scientific | Cat #A12379 |
| Rabbit anti TRIM46 | C. Hoogenraad | (17) |
| Rabbit anti-GFP | MBL International | Cat #598 |
| Rabbit anti-Laminin antibody | Abcam | Cat #ab11575 |
| Rat Anti-Tubulin antibody [YL1/2] | Abcam | Cat #ab6160 |

#### Table of DNA constructs

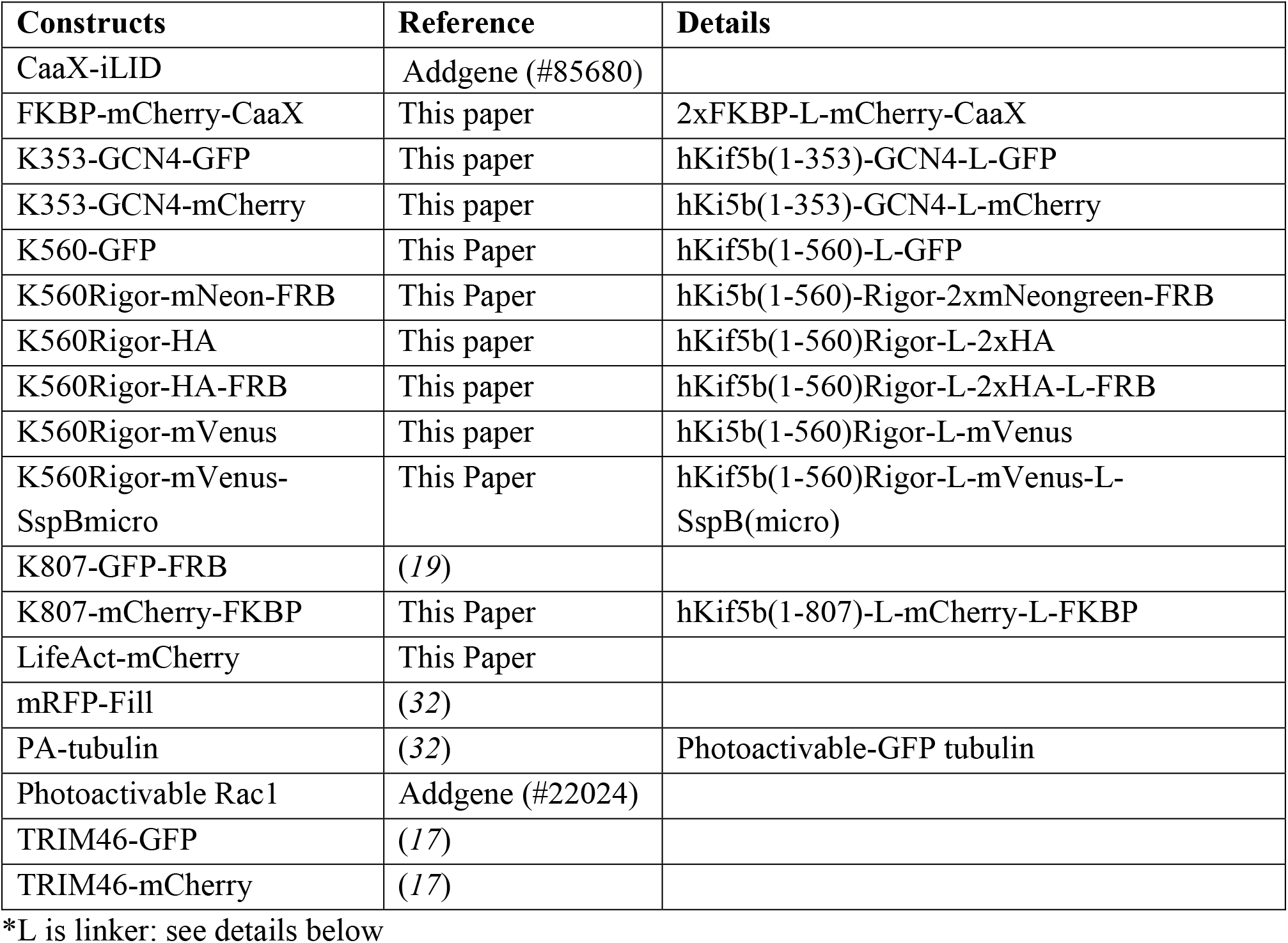

| **Constructs** | **Reference** | **Details** |
| --- | --- | --- |
| CaaX-iLID | Addgene (#85680) |  |
| FKBP-mCherry-CaaX | This paper | 2xFKBP-L-mCherry-CaaX |
| K353-GCN4-GFP | This paper | hKif5b(1-353)-GCN4-L-GFP |
| K353-GCN4-mCherry | This paper | hKi5b(1-353)-GCN4-L-mCherry |
| K560-GFP | This Paper | hKif5b(1-560)-L-GFP |
| K560Rigor-mNeon-FRB | This Paper | hKi5b(1-560)-Rigor-2xmNeongreen-FRB |
| K560Rigor-HA | This paper | hKif5b(1-560)Rigor-L-2xHA |
| K560Rigor-HA-FRB | This paper | hKif5b(1-560)Rigor-L-2xHA-L-FRB |
| K560Rigor-mVenus | This paper | hKi5b(1-560)Rigor-L-mVenus |
| K560Rigor-mVenus-SspBmicro | This Paper | hKif5b(1-560)Rigor-L-mVenus-L-SspB(micro) |
| K807-GFP-FRB | (19) |  |
| K807-mCherry-FKBP | This Paper | hKif5b(1-807)-L-mCherry-L-FKBP |
| LifeAct-mCherry | This Paper |  |
| mRFP-Fill | (32) |  |
| PA-tubulin | (32) | Photoactivable-GFP tubulin |
| Photoactivable Rac1 | Addgene (#22024) |  |
| TRIM46-GFP | (17) |  |
| TRIM46-mCherry | (17) |  |

\*L is linker: see details below

#### DNA constructs

All constructs used in this study, as well as their source, are listed in the table of DNA constructs. New constructs generated for this study were cloned into the mammalian expression vector pβactin-16-pl (chicken β-actin promoter) (*33*) by conventional molecular cloning and PCR-based cloning. For all motor constructs, the human Kif5b sequence (P33176) was used. To avoid dimerization of the motile kinesin constructs with the K560Rigor, we replaced the dimerization domain of Kinesin-1 with a GCN4 leucine zipper (LEDKIEELLSKIYHLENEIARLKKLIGEI) for the K353-GCN4-GFP and K353-GCN4-mCherry constructs (*34*). Synthetic, 29 amino acid GGGS linkers were added between domains as indicated by ‘L’ in the details of the constructs in the table above (*34*). To anchor microtubules to the plasma membrane, we introduced the G234A mutation in hKif5b to create K560Rigor (*35*). The sequence for mNeongreen and the linker sequences flanking this fluorophore were provided by Allele Biotechnology (*36*). mVenus and SspB(micro) domains were derived from pB80-KIF1A(1-365)-mVenus-SspB(micro)(*34*). The CAAX motif CVIM was derived from Venus-iLID-CaaX (Addgene #60411). The FKBP-FRB heterodimerization modules were described previously in (*19*).

#### Animals, neuron culture and transfection

All animal experiments were approved by the Dutch Animal Experiments Committee (DEC, Dier Experimenten Commissie) (license number AVD1080020173404) and were in line with the institutional guidelines of Utrecht University. All animal experiments were performed in compliance with Dutch law (Wet op de Dierproeven, 1996) and European regulations (Directive 2010/63/EU). Pregnant Wister rats (Janvier), which were not involved in any previous experiments and were ≥10 weeks of age, were used in this study. For the primary neuron culture, cells were derived from hippocampi of embryonic day 18 (E18) pups (both genders). Hippocampi were dissociated using a method involving both enzymatic and mechanical dissociation (*37*). After dissociation, neurons were first electroporated (see below) or plated directly at a density of 50,000 cells per well in a 12-well plate on coverslips coated with poly-L-lysine (37.5 µg/mL, Sigma Aldrich) and laminin (1.25 µg/mL, Roche). The primary cultures were maintained in Neurobasal medium (NB, Gibco) supplemented with 2% B27 (Gibco), 0.5 mM glutamine (Gibco), 15.6 µM glutamate (Sigma-Aldrich) and 1% penicillin/streptomycin (Gibco) (from now on referred to as full medium) at 37°C and 5% CO_2_.

#### Electroporation and transfection

For imaging DIV1-3 neurons electroporation was preformed whereas for neurons >DIV3, lipofectamine transfection was used. Electroporation of hippocampal neurons was performed directly after dissociation. The cell suspension was spun down at 200g for 5min, supernatant was removed and neurons were resuspended carefully in electroporation buffer (12.5 mM NaCl, 123 mM KCl, 20 mM KOH, 10 mM EGTA, 4.5 mM MgCl_2_, 20 mM PIPES, pH 7.2) mixed with 20% FCS. The electroporation buffer and FCS aliquots were stored at −20°C and mixed after being warming up separately. Vigorous resuspension using a pipette can damage the neurons and small clumps of neurons that did not resuspend after 4-5 times mixing with the pipette were removed from the solution whenever possible. The resuspended neurons were mixed with 1-3 µg plasmid DNA and placed in electroporation cuvette (density of 200k-1 million neurons/cuvette) (Bio-Rad GenePulser, 0.2cm gap) and electroporated in a Nucleofector 2b device (Lonza) on program O-003/rat hippocampal neurons. After electroporation, neurons were diluted in full medium and divided over multiple wells at ∼50,000 neurons per 18 mm coverslip, which were coated as described above. For DIV1 neuron imaging, the following amounts of DNA were used: K353/K560 (1.5-2 µg), TRIM46 (0.8 µg), K560Rigor (0.3 µg), iLID-CaaX (1 µg), PA-tubulin (2 µg), mRFP-fill (0.3 µg), Photoactivable Rac1 (1 µg). For Lipofectamine transfection, 50k neurons per well of a 12-well plate were directly plated in full medium onto 18-mm coverslips coated as described above and transfected at the indicated times using Lipofectamine 2000 (Invitrogen). For one well of a 12-well plate, 1.8 μg DNA was mixed with 3.3 μl Lipofectamine in 200μl NB and incubated for 30 min at room temperature. Meanwhile, conditioned medium, the medium in which neurons were grown, was transferred to a new plate, and transfection medium (NB medium supplemented with 0.5 mM glutamine) was added. The DNA/lipofectamine mix was added to the neurons in transfection medium and incubated for 45 min at 37°C and 5% CO_2_. Neurons were rinsed by dipping coverslips in pre-warmed NB medium and placed back into conditioned full medium for 1-2 days prior to imaging.

U2OS cells were purchased from ATCC and cultured in DMEM medium containing 10% FCS and 50 µg/ml Penicillin/Streptomycin at 37°C and 5% CO_2_. Cells were confirmed to be free of mycoplasma. U2OS cells were plated on 25mm diameter coverslips 2 days prior to transfection. For transfection of one 6 well with a 25 mm coverslip, 6μl of Fugene6 transfection reagent (Promega) was added to 100μl of Leibovitz’s L-15 medium followed by addition 0.3μg - 2μg of DNA. The mix was incubated for 5 min at room temperature and then homogeneously added to the medium in a well.

Developmental stages of neurons were categorized as stage 2, stage 3 and stage 4 by morphology (*5*). Days in vitro (DIV) were counted after seeding the dissociated neurons: DIV1-2 (26-48 h), DIV2-3 (48-72 h) and DIV7-8 (168-192 h) neurons were used for stage 2, stage 3 and stage 4 of neuron development, respectively (*5, 9*).

#### Polyacrylamide hydrogel stiffness gradients

The hydrogel stiffness gradients for culturing embryonic rat hippocampal neurons were based on polyacrylamide. We combined elements from other approaches (*4, 11, 38*) to optimize the fabrication of hydrogel gradients. First, coverslips were silanized according to method described previously(*38*). Briefly, 18 mm glass coverslips were plasma treated using a plasma cleaner (Harrick Plasma Cleaner PDC-002) for 3 min and immediately treated with 2%(v/v) 3-(trimethoxysilyl)propyl methacrylate and 1% (v/v) acetic acid in ethanol for 10min in a beaker with occasional manual shaking. Coverslips were washed twice with ethanol and then air-dried using coverslip holders. Dried coverslips were baked at 120°C for 2h and stored in dry place at room temperature up to a month. To generate polyacrylamide hydrogel gradients, two acrylamide solutions, soft (0.4 kPa) and stiff (7 kPa) were prepared to create stiffness gradient of 0.4-7 kPa. Gel pre-mixes were prepared by mixing acrylamide and bis-acrylamide solutions. The desired Young’s modulus of the pre-mixes was adjusted by mixing pre-defined ratios of 40% (w/v) acrylamide and 2% (w/v) N,N-methyl-bis-acrylamide cross-linker in PBS. Gel premix solutions (200μl) corresponding to 0.4 kPa (5% AA and 0.04% Bis-AA) and 7 kPa (12% AA and 0.03% Bis-AA) were prepared. Fluorescently labelled 0.2 μm carboxylated beads (dark red fluorescent (660/680) were added at 3.6*10^9^ beads per 100μl to the stiff pre-mix solution. Bead solution was vortexed for 3 min prior to adding to the gel solution. Silanized coverslips were placed onto parafilm. Polymerization was initiated by adding 2 μl of 10% ammonium persulfate and 1μl of *N,N,N’,N’*-tetramethylethylenediamine (TEMED) to the solutions. 8 μl droplets of each stiff and soft premixes were put onto the silanized coverslip 2-3 mm apart on one side of the 18 mm coverslip. A 14 mm glass coverslip was placed onto the droplets by gently dropping it from one end to the other end of 18-mm coverslip, leading to in situ mixing of acrylamide solution by diffusion(*11*). Gels were polymerized for 45-50 minutes at room temperature and then covered with 2 ml of PBS for 30 min to facilitate detachment of gels from coverslips. Then, the 14 mm coverslip was gently detached using needle and tweezers. Acrylamide hydrogels attached strongly to the silanized coverslips and were placed in PBS solution at 4°C up to 4-5 days. To allow Laminin to adhere to hydrogel substrates, gels were incubated in hydrazine hydrate for 4h, 5% acetic acid for 1h and thoroughly washed with PBS 3 times. Gels were incubated in 37.5μg/ml poly-l-lysine in PBS for 2 days and then incubated with 12.5μg/ml Laminin in PBS for 2h before plating the neurons. To measure the coating of Laminin on hydrogel gradient substrates, gels were incubated with rabbit anti-Laminin antibody diluted 1:200 in 3% bovine serum albumin (BSA) in PBS. Gels were washed three times with PBS and then incubated with anti-rabbit Alexa488 antibody for 1h followed by three washes with PBS. Gels were mounted onto glass slides using ProLong diamond mountant. Images of beads and Laminin were acquired every 300 μm across the gradient using 40x objective of Nikon spinning disk microscope (see below).

A control experiment was done to test whether the axon formation is sensitive to gradient of fluorescent beads (Fig S3). 3 kPa rigidity was chosen as an intermediate rigidity of the 0.4-6 kPa gradient for the control experiment. Two polyacrylamide premixes corresponding to 3 kPa (7.5% AA and 0.06% Bis-AA) were prepared. Fluorescent beads were added to one solution and the gradient was prepared as described above. Bead gradient was analyzed and axon orientation was determined as described below in image analysis section.

#### Rheology measurements

Mechanical properties of the hydrogels were assessed by oscillatory shear measurements, using a DHR-2 rheometer (TA Instruments), with a plate-plate geometry (20 mm diameter) and Peltier hood to prevent evaporation. The viscoelastic linear regime (LVR) was determined for all formulations by an amplitude sweep test at angular frequency *ω*= 10 rad/s. Hydrogel formation was monitored by a time sweep test (oscillation strain *γ*= 2%, *ω*= 10 rad/s, 30-40 minutes), and frequency-dependent measurements were carried out in the frequency range 0.1-100 rad/s, at a strain of *γ*= 2% (safely within LVR). All measurements were conducted at 20 ^0^C and in triplicates, unless noted otherwise. The stiffness of the gels was taken as the value of the storage modulus G’(*ω*) at *ω*= 1 rad/s, as measured by the frequency sweep test. The values are reported in Fig. S2E.

#### Live-imaging assays

For live-cell imaging, 18 mm coverslips were mounted in a Ludin chamber (Life Imaging Services) covered with 1 ml of conditioned medium. The chamber was mounted in Tokai hit chamber maintained at 37°C and supplied with 5% CO2. Cells were imaged using Nikon Spinning Disk Confocal microscope with following configuration: Nikon Eclipse Ti (Nikon) with Perfect focus system (Nikon) equipped with Photometrics prime BSI camera or Evolve Delta 512 EMCCD camera (Photometrics), Plan Apo 60x N.A. 1.4 oil or Plan Fluor 40x N.A. 1.30 oil objectives (both Nikon), Vortran Stradus 405nm (100mW), Cobolt Calypso 491nm (100mw) and Cobolt Jive 561 nm (100mW) laser, ASI motorized stage MS-2000-XYZ with Piezo top plate, ET-mCherry (49008), ET-GFP(49002), ET-GFPmCherry(59022) filters (all Chroma), ILAS2 FRAP module (Roper scientific, now Gataca systems), Spinning disk-based confocal scanner unit (CSU-X1-A1, Yokogawa) and MetaMorph 7.7 software (Molecular Devices).

The electroporation protocol caused death of a small proportion of neurons within 12-24 h after electroporation. Hence, neuron morphology was checked with transmitted light prior to live imaging. Healthy neurons with low to moderate expression level of protein were carefully chosen for live-cell imaging to avoid overexpression artifacts and phototoxicity. To further reduce phototoxicity, low laser powers (4-10%) were used. Illumination time between 200-300ms and an EM gain of 900-950 were used with EMCCD camera.

#### Photoactivation of Rac1 and PA-tubulin

For Rac1 photoactivation, neurons were electroporated with photoactivable Rac1 construct and human Kinesin-1(Kif5b(1-353)-GCN4-mcherry). DIV1 neurons expressing low levels of Kinesin-1 were selected for imaging and a Region of interest (ROI) was drown around a neurite. Neurons were imaged with 561nm in the ‘Dark’ meaning without 405 nm for 15-40 min prior to photoactivation. Using a custom-made journal in MetaMorph software, 405nm laser was used to illuminate the region of interest every 5-10 seconds for total of 30-40 min. Kinesin-1 was imaged with 1 frame per minute. For the photoactivatable tubulin experiments, neurons were electroporated with PA-tubulin and mCherry-fill constructs and imaged at stage 2 or stage 3. For imaging of stage 4 neurons, neurons were transfected with Lipofectamine with the same constructs and imaged 24-48 h after transfection. Using the ILAS2 FRAP module, single or multiple regions of interests of 10-15µm length were drawn along stage 2 neurites (for stage 2 neurons) or axons and dendrites (stage 3 and 4 neurons). All the ROIs on a neuron were illuminated at once with 20-27% laser power of 405 nm Cobolt Calypso (100mW) laser with 5-6 repetition. Upon photoactivation, the PA-tubulin signal was imaged using 491nm laser excitation for 30-60 minutes with 1 frame per minute.

#### Microtubule anchoring assays

For Rapalog-induced microtubule anchoring, neurons were electroporated with FKBP-mCherry-CaaX as a membrane anchor, K560Rigor-HA-FRB as a microtubule anchor and K353-GCN4-mCherry for monitoring Kinesin-1 transport. DIV1 stage 2 neurons with low expression levels of Kinesin-1 were selected. Rapalog was added at a concentration of 200nM to anchor the Kinesin-1 rigor-decorated microtubules to the cell membrane. As a control experiment, the experiment was repeated using K560Rigor-HA without the FRB domain (Fig. S10). Cells were either imaged live or after fixation with pre-warmed 4% paraformaldehyde in PBS after 2 hours of Rapalog addition.

For light-induced microtubule anchoring, neurons were electroporated with iLID-CaaX as a membrane anchor, K560Rigor-mVenus-SspB as a microtubule anchor and K353-GCN4-mCherry for monitoring Kinesin-1 transport. Neurons were imaged on the Nikon spinning disk microscope described above, using the 40x oil objective and the ILAS2 FRAP module for photoactivation. Neurons were protected from stray light prior to imaging by wrapping the 12-well plate in aluminum foil. The microscope set up was surrounded by black curtains to avoid any stray light activating the photo-sensitive module. DIV1 stage 2 neurons with low expression level of Kinesin-1 were selected for imaging. Kinesin-1 (Kif5b-mcherry) was imaged with 561nm laser every 60 seconds, while local illumination using a 405 nm laser was performed every 5-10 seconds to achieve light-induced anchoring. As a control experiment, neurons were electroporated with iLID-CaaX and K560Rigor-mVenus without the SspB domain for light-induced microtubule anchoring (Fig. S10).

#### Pharmacological treatments and immunostainings

Neurons were electroporated with a Kinesin-1 construct and plated onto coated 18 mm coverslips. Thirty hours after plating, Rac inhibitor EHT1864 (TOCRIS #3872) or Cdc42 inhibitor ML141 (TOCRIS #4266) was added to the cells at a concentration of 20 µM for 2h and then fixed for immunostaining. DMSO or distilled water was added as control. DIV1 neurons were fixed in pre-warmed (37°C) 4% Paraformaldehyde and 0.1% gluteraldehyde in PBS for 15 min. After 3 washes with PBS, cells were permeabilized with 0.2% triton X-100 in PBS for 10 min and then washed 3 times with PBS. Cells were blocked for 40-60 min with 3% BSA in PBS followed by 1h incubation in primary antibodies diluted in 3% BSA. After 3 washes with PBS, cells were incubated with secondary antibodies diluted in 3% BSA for 30min. After 3 washes with PBS coverslips were mounted in proLong Diamond antifade mountant (Thermo Fischer Scientific).

#### Analysis of stiffness gradients

Images of neurons on gradient gels with fluorescent beads were imaged using a 40x objective on the spinning disk microscope described above. Single, isolated neurons were imaged within 200×150µm regions by acquiring a z-stack with 1μ m z-plane distance covering 12-15µm distance. To analyze the rigidity along the growing neurites or axon, multi-channel images comprising of bead channel and Kinesin-1 or Tau staining were acquired. A maximum intensity *z-*projection was performed for all channels using FIJI. Beads were counted using the <Find maxima> plugin in FIJI within a 300×300 pixel (∼45 x45 µm) grid area of the bead image. The value of prominence in the plugin was carefully adjusted for each experiment to obtain detection of all beads in the region of interest and to avoid false positive and false negative detection of beads.

To derive the relation between the number of beads and local gel stiffness, a gel consisting of only the stiffer, bead-containing solution was made. Images of the beads were taken in similar fashion as described above, resulting in an estimate for the number of beads per 10,000 µm^2^ for the stiffest rigidity of 7 kPa, whereas the softer gel (0.4 kPa) featured density of 0 beads per 10,000 µm^2^. Based on previous work (*11*), a linear relationship between the number of beads and stiffness within 0.4-7 kPa rigidity range was assumed, resulting in the following calibration scale: *y*= 0.44 + 0.001987\**x*, where y is stiffness in kPa and x is number of beads per 10,000 µm^2^.

Because beads were counted in grid areas of ∼45 µm x 45 µm, an interpolation method was used to better estimate local gradients. The interpolation method (image Resize option, bilinear interpolation) in FIJI performs interpolation, but places the original pixel values in the top left of each interpolated region. To correct for this shift, we performed ‘Matrix padding’ and then cropped the image so that the value of the bead count was at the center of each grid unit of 45 µm x 45 µm in the interpolated image (Fig S2). Only region of interest that featured uniform bead gradients were chosen for analysis. Although the global gradient of the gels was known, for each analyzed neuron we determined the exact local direction of the gradient. To this end, a 80 µm radius circle was drawn centered around the cell body and the maximum difference in stiffness was determined between for 18 equidistant pairs of points on this circles. The points corresponding to the maximum difference defined the axis against which the angles of axons, dendrites and immature neurites were measured. Neurites with angles within the range of < 90° and > −90° were counted as facing toward the softer matrix, whereas neurites with angles within the range of > 90° and < −90° are counted as facing towards the stiffer matrix.

#### Analysis of live-imaging assays

Time-lapse TIFF images were processed in FIJI. Intensities of the labelled proteins were measured by drawing ROIs enclosing different neurites or the soma. Background subtraction was performed by subtracting the mean background intensity multiplied by the ROI area from the raw intensity of the ROI for every frame. In some cases where ROI intensities were very low, the mean value of the background intensity was subtracted to avoid negative intensity values. For generating movies, the average background intensity was subtracted from the entire stack of images and a Gaussian blur of 1 pixel was applied. For the TRIM46 intensity analysis in Fig. S7, the TRIM46 intensity in each neurite was measured at every time frame for a total of 2 hours. Then, the TRIM46 intensity at each frame was divided by the minimum TRIM46 intensity in a neurite during those 2 hours to obtain the fold change of intensity at every frame and the maximum fold change for each neurite was plotted. For quantification of Kinesin-1 intensity, the intensity in each neurite was measured at every time frame and divided by the total Kinesin-1 intensity within all neurites to obtain the percentage of Kinesin-1 accumulation in a neurite at every time frame.

To analyze microtubule network movement using PA-tubulin, the time-lapse movies of each photoactivated region along axon, dendrites or stage 2 neurites were converted into a kymograph. To generate a kymograph, a segmented line of 3.5 µm thickness, starting closer to the cell body was drawn along the photoactivated-GFP tubulin regions. The ‘KymoResliceWide’ plugin in FIJI was used to obtain the kymographs. Negative and positive velocities were assigned to the retrograde (towards the cell body) and anterograde (away from the cell body) movements, respectively.

#### Statistical analysis

Data sets were tested for normal (Gaussian) distribution using the Shapiro-Wilk normality test. In case of normal distribution, statistical comparisons between two groups were done using two-tailed Student’s t test. If data did not follow a normal distribution, a Mann-Whitney non-parametric test was used to compare the differences between different conditions. To compare proportions of different outcomes between two or more conditions, the Chi-square test used. To compare difference between values before and after treatment for the same sample, the non-parametric Wilcoxon matched-pair test was used. Data was plotted and statistical tests were performed using GraphPad Prism9. Error bars indicate standard deviations.

## SUPPLEMENTAL FIGURES

**Figure S1.**
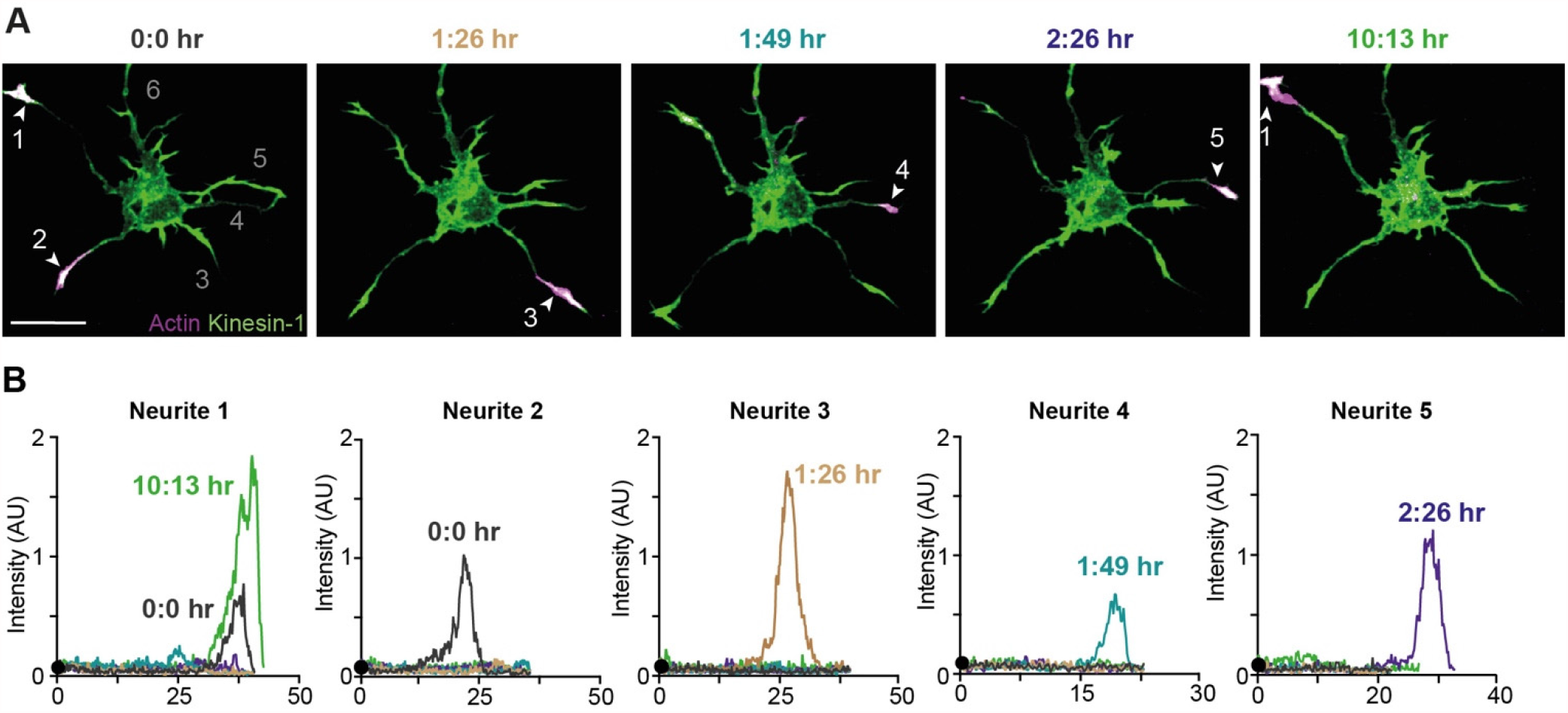
Transient accumulation of Kinesin-1 in neurites of stage 2 neurons grown on glass. (A) Selected time-frames of a stage 2 neuron expressing Kinesin-1 (magenta) and Lifeact (Green). White arrowhead indicates neurites with Kinesin-1 accumulation. (B) Intensity profiles along the length of neurites 1-5 for the different time frames shown in (A). Colors are assigned to different time frames. Black dot represents the position of cell body. a.u., arbitrary units.

**Figure S2.**
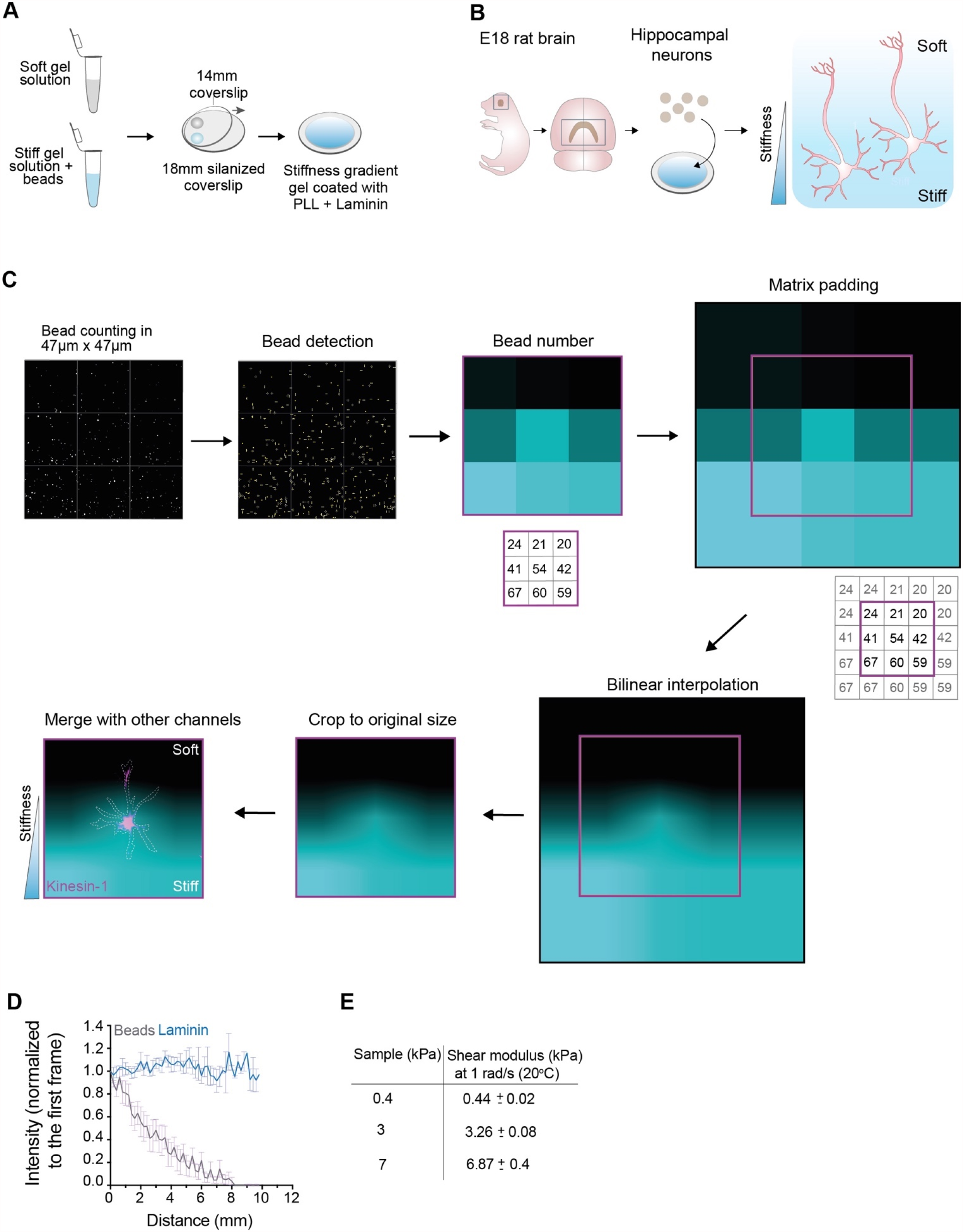
Fabrication of elasticity gradients using poly-acrylamide hydrogels. (A) Pre-mixes of gel solutions for stiff and soft gels solutions were prepared and from each solution one drop of 8 μl was placed onto a silanized glass coverslips. A 14 mm glass coverslip was placed onto the droplets by gently dropping it from one end to the other end, followed by polymerization. After polymerization, gels were coated with poly-L-lysine (PLL) and Laminin. (B) Hippocampal neurons from embryonic Rat brain (E18 stage) were dissociated and seeded onto the PLL and Laminin coated hydrogels gradients. Neuron growth was tracked on different days to capture different stages of development. (C) The elasticity gradient of acrylamide hydrogels was derived from images of fluorescent beads embedded within the gel. Beads were detected in squares of fixed size. In this example, a 3×3 pixel image was generated using the bead counts within the 9 squares. Before interpolation, extra rows and columns were added at the edges of the matrix (numbers in grey, see methods), generating a 5×5 matrix. This 5×5 pixel image was scaled up using bilinear interpolation, followed by cropping to the original size of the image (magenta box). The cropping area was positioned so that the original value of the bead count in the grid was at the center of each grid area in the interpolated image. The final cropped gradient image was merged with other imaging channels. (D) Quantification of the bead density and Laminin intensity on the stiffness gradient hydrogels (n=11). Error bars represent SEM. (E) Shear modulus values of gel premixes of different stiffnesses. The premixes for 3 and 7 kPa gels contained beads.

**Figure S3.**
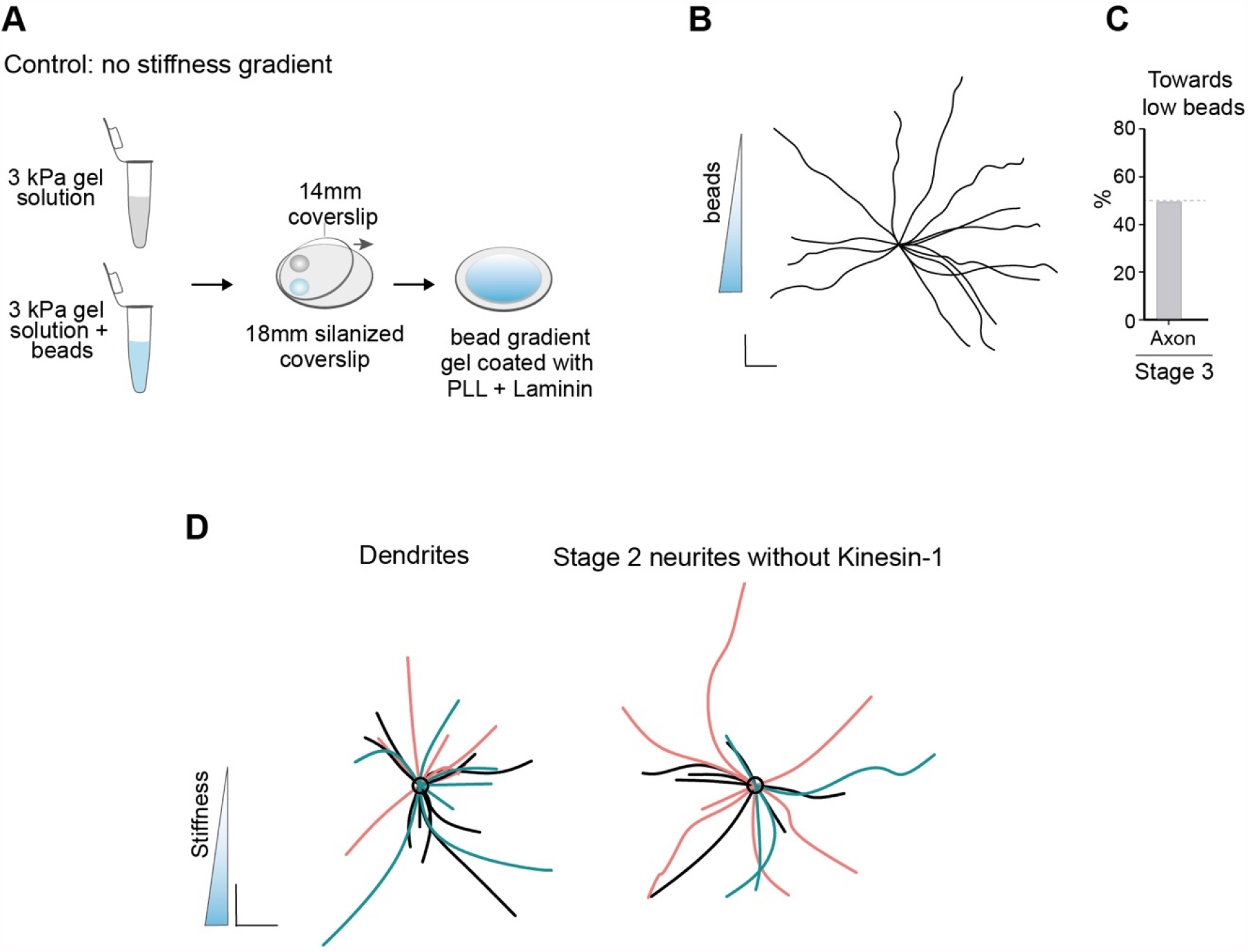
Fluorescent bead gradient does not affect axon formation. (A) A bead gradient was generated by mixing a solution for 3 kPa containing fluorescent beads with a similar solution without beads. (B) Traces of stage 3 axons on 3 kPa polyacrylamide gels with a bead gradient show no bias for the direction of the gradient. Scale bar 20 µm. (C) Quantification of percentage of axons growing towards lower bead density. (D) Orientations of dendrites (n=25) and stage 2 neurites (n=18) on elasticity gradients from 3 independent experiments. Different colors represent traces from independent experiments.

**Figure S4.**
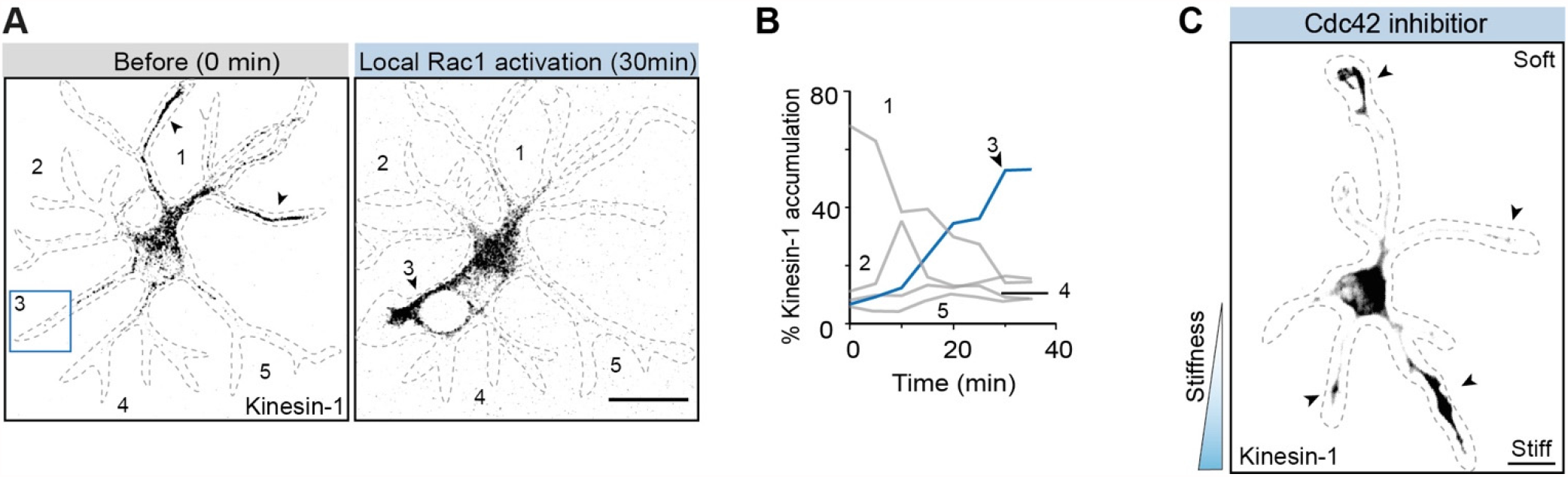
Local Rho-GTPase activity and selective Kinesin-1 accumulation. (A) Rac1 photoactivation in stage 2 neuron redirects Kinesin-1(K345-GCN4) accumulation from neurite 1 to neurite 3. Scale bar 10 µm. (B) Time traces of Kinesin-1 accumulation in different neurites show an increase in neurite 3 upon local Rac1 activation. (C) Loss of selective transport to a neurite upon Cdc42 inhibitor in a neuron grown on elasticity gradient. Scale bar, 10 µm.

**Figure S5.**
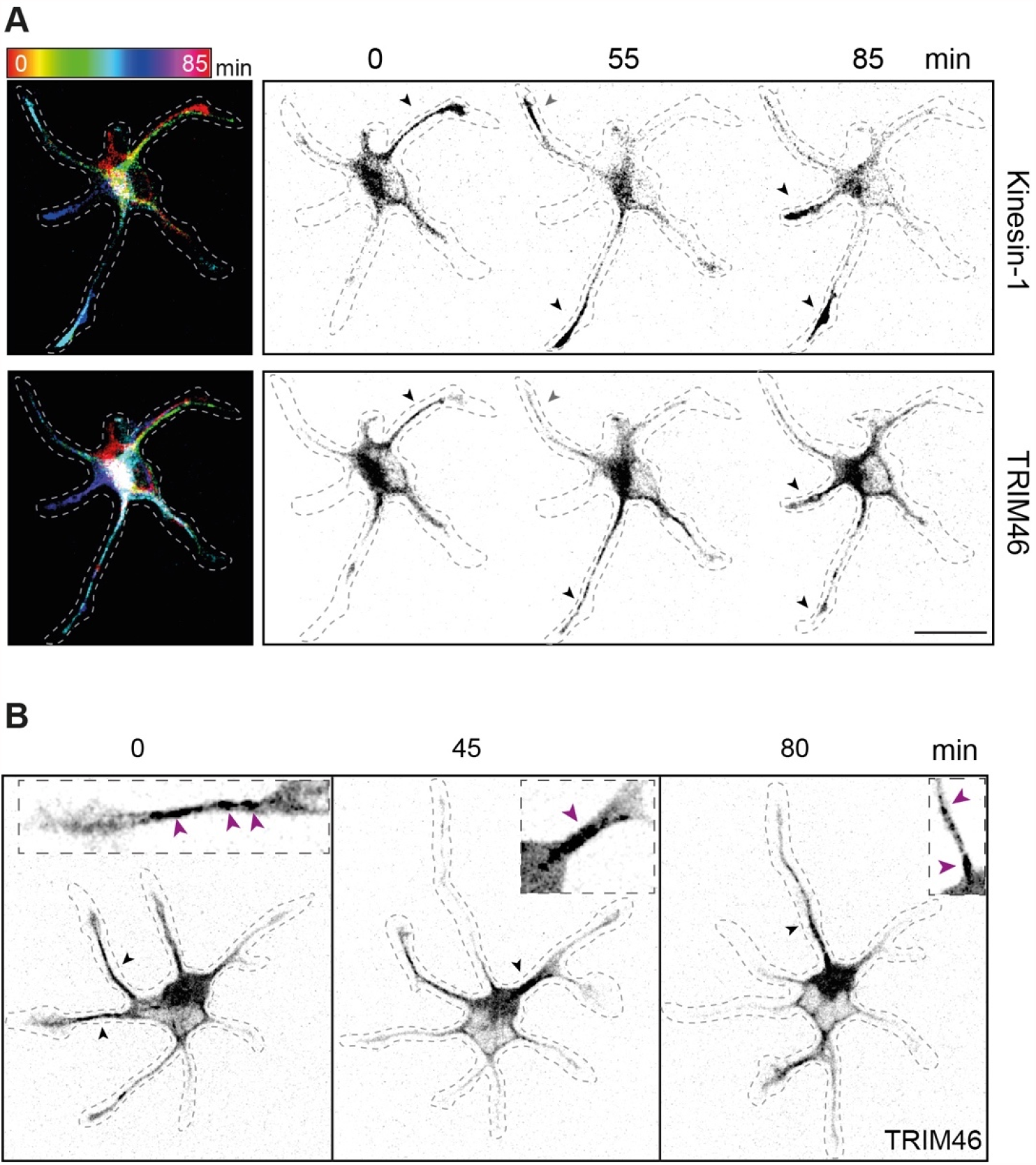
Correlation of Kinesin-1 and TRIM46 accumulation in stage 2 neurons. (A) Left, time-color coded images of Kinesin-1 and TRIM46 in the stage 2 neuron depicted in Fig 3, A. Inverted grey scale images of Kinesin-1 and TRIM46 at different time-frames show that Kinesin-1 and TRIM46 accumulate in same neurites as indicated by the black arrowheads. The grey arrowhead marks an example where TRIM46 loss from a neurite precedes Kinesin-1 loss. (B) Time frames of stage 2 neurons expressing TRIM46 and mRFP-fill and grown on glass. Inverted grey scale images show TRIM46 patches appearing in alternating neurites over time, marked by black arrowheads. Insets show zoom-in images of TRIM46-enriched areas in neurites. TRIM46 patches (magenta arrowheads) are enriched at the base and proximal part of the neurites.

**Figure S6.**
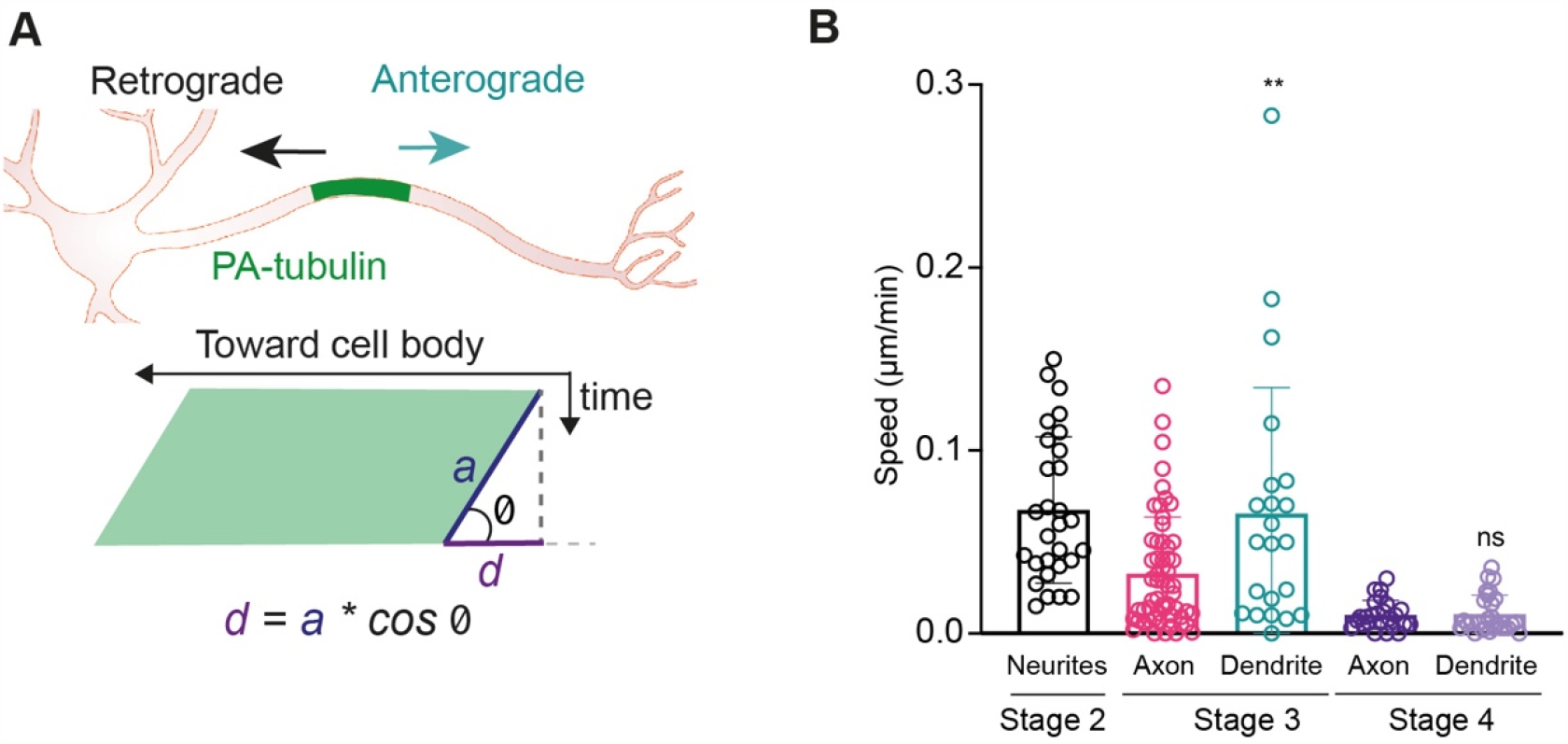
Microtubule network mobility is reduced in stage 3 axon. (A) Illustration of kymograph analysis used for Fig. 3. (B) Quantification of absolute speeds of microtubule network movements (from data in Fig 3H). In stage 3 axon the mobility is lower than in dendrites. **≤0.01 (Two tailed student’s t test).

**Figure S7.**
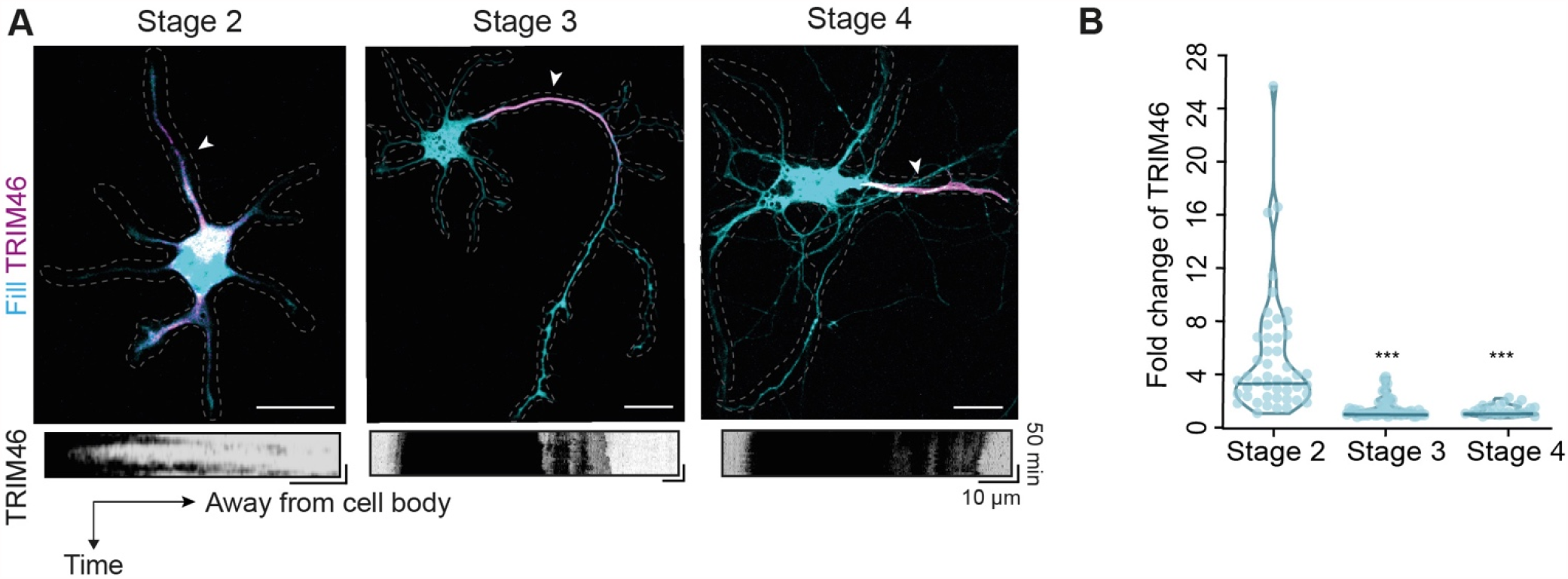
TRIM46 fluctuations between neurites reduce during development. (A) Top: snapshots of stage 2-4 neurons expressing TRIM46 and mRFP-fill. Bottom: inverted grey scale images show kymographs of TRIM46 within the neurites marked by white arrowheads on top images. Scale bar 10 µm (*x*) and 50 min (*y*). (B) Fold change of TRIM46 intensity in a neurite with respect to the minimum intensity value over the course 2 h. stage 3 and 4 values were statistically compared with stage 2. ***P≤0.001 (Mann-Whitney test).

**Figure S8.**
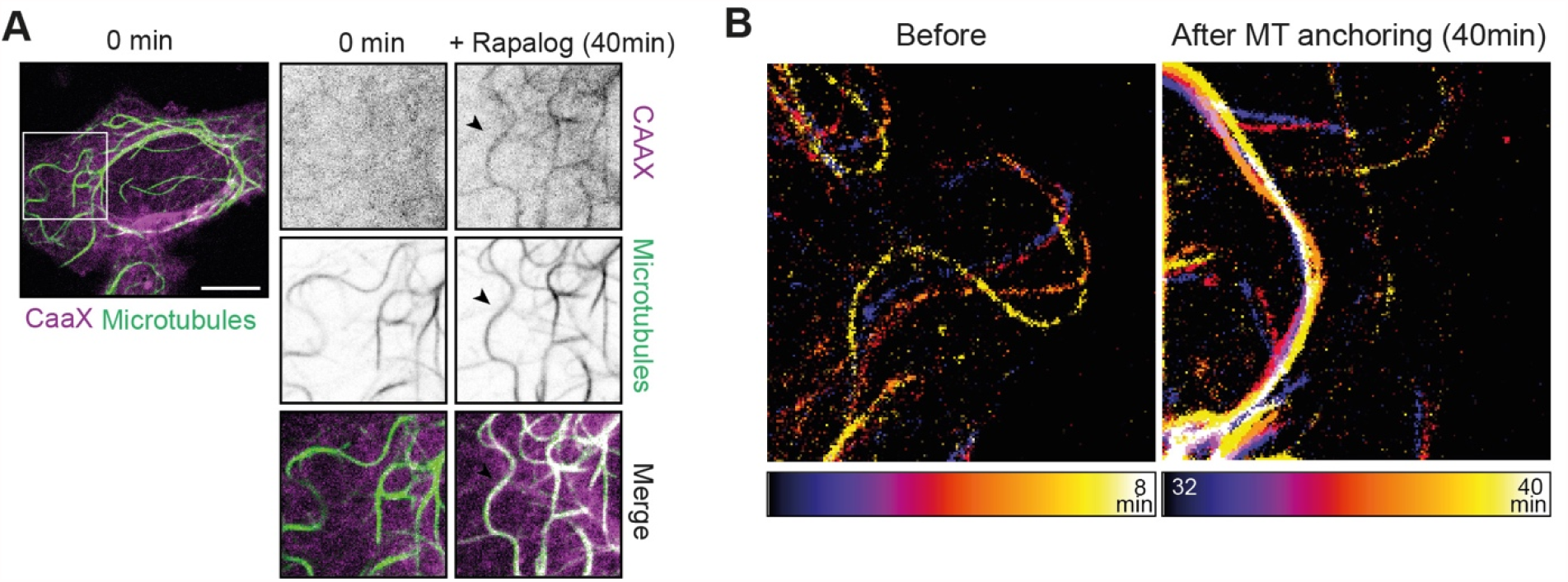
Transient accumulation of TRIM46 in neurites reduces during development. (A) Left: U2OS cell expressing CaaX-FKBP and microtubules decorated with the FRB-microtubule anchor (Kinesin-1-rigor) before Rapalog addition (time 0 min). Right: zooms of the area in rectangle on left image. Individual images of CaaX, FRB-microtubule anchor, and merge, before (0 min) and after Rapalog addition (40 min). Black arrowhead shows appearance of microtubule morphology in the CaaX channel after Rapalog addition. Scale bar, 10 µm. (B) Time-color coded overlays from a time-lapse recording demonstrating reduced microtubule mobility after Rapalog addition.

**Figure S9.**
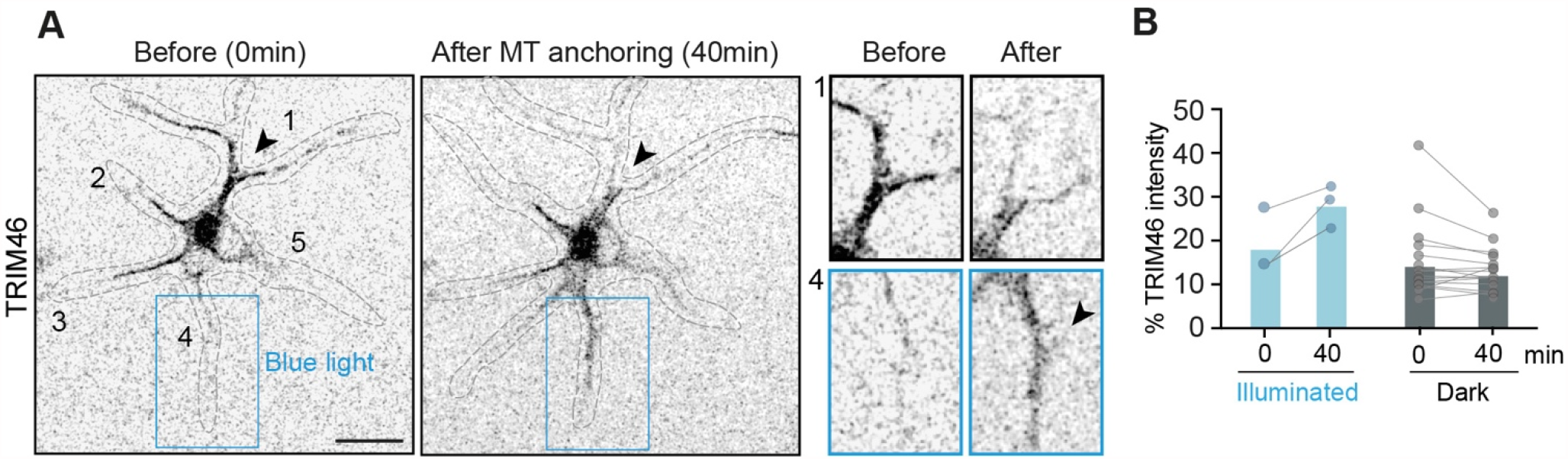
Local anchoring of microtubules drives selective TRIM46 accumulation. (A) Snapshots of a stage 2 neuron expressing iLID-CaaX, SspB-microtubule anchor and TRIM46. Upon blue light illumination, TRIM46 is relocated from neurite 1 at 0 min to neurite 4 at 40 min. Blue rectangle represents area illuminated with blue light. Black arrow marks area with reduction of TRIM46 intensity from neurite 1 upon microtubule anchoring. On the right, zoom-in images of neurite 1 and 4 showing loss of TRIM46 patches from neurite 1 and increase in TRIM46 intensity in neurite 4, marked by arrowhead. Scale bar, 10 µm. (B) Quantification of percent TRIM46 accumulation in neurites activated with blue light for 40 min (blue dots) (n=3 pairs) and non-illuminated neurites (grey dots) (n=16 pairs) Rectangles represent median values.

**Figure S10.**
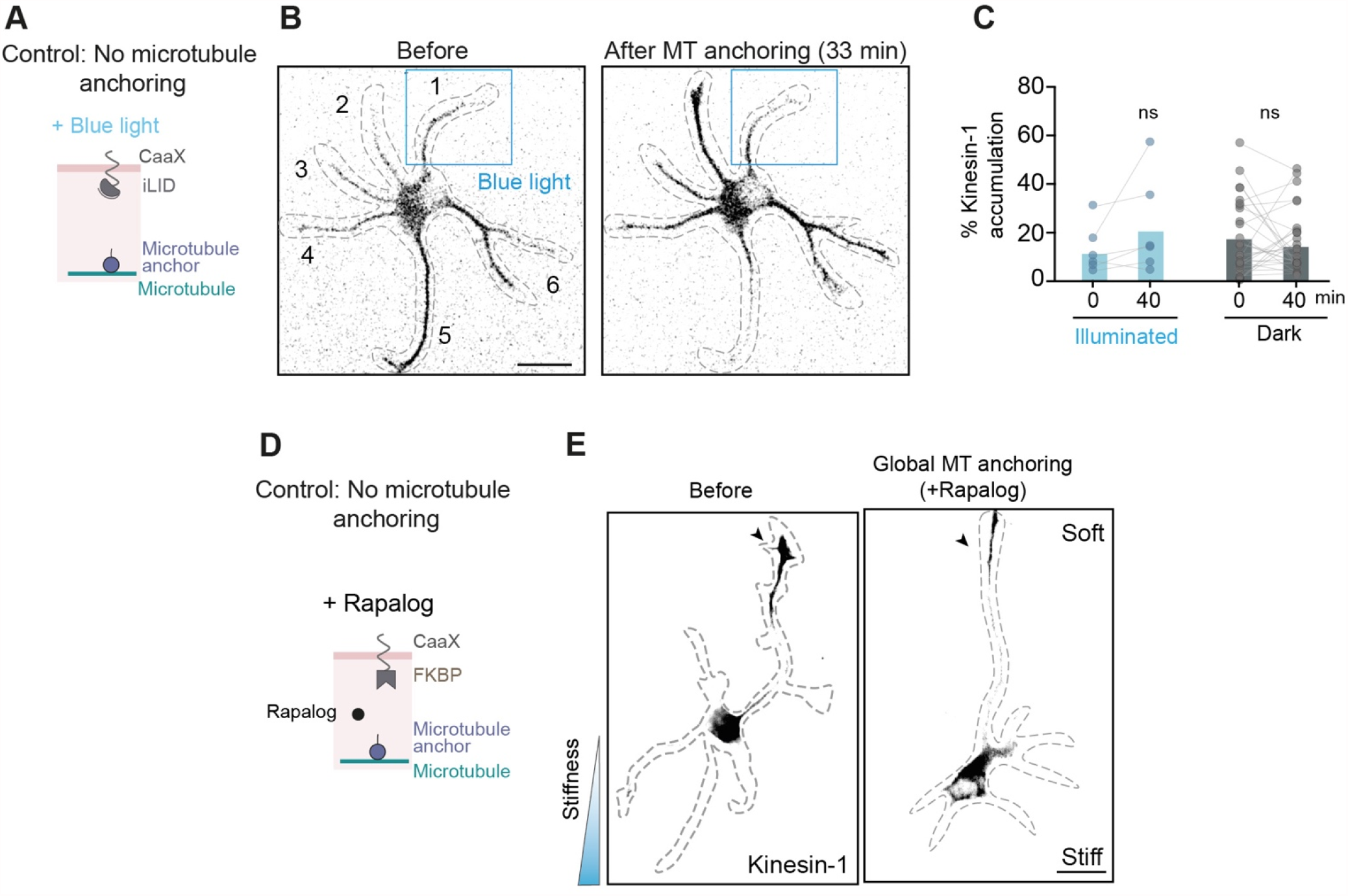
Control experiments for the global and local microtubule anchoring assays. (A) Control experiment for light-induced local microtubule anchoring assay. The blue light illumination cannot induce binding of iLID-CaaX to microtubule anchor without the SspB domain. Consequently, microtubules do not get anchored to the plasma membrane after blue light illumination. (B) Immunofluorescence of stage 2 neuron growing on glass and expressing iLID-CaaX, microtubule anchor and Kinesin-1. Blue rectangle represents area that was illuminated with blue light. Kinesin-1 accumulation changes from neurite 5 and 6 to 2 and 6 after microtubule anchoring but does not enrich into neurite 1. (C) Quantification of percent Kinesin-1 accumulation in neurites activated with blue light for 40 min (blue dots) (n=6 pairs) and non-illuminated neurites (grey dots) (n=28 pairs). Rectangles represent median values. ns: non-significant (two-tailed Wilcoxon matched-pairs test). (D) Control experiment for Rapalog-induced global microtubule anchoring assay. Rapalog addition cannot induce binding of FKBP-CaaX to microtubule anchor without the FRB domain. Consequently, microtubules do not get anchored to the plasma membrane after Rapalog addition. (E) Immunofluorescence of stage 2 neuron growing on stiffness gradient and expressing FKBP-CaaX, microtubule anchor and Kinesin-1. Before and after Rapalog addition, the neurite with selective Kinesin-1 accumulation is oriented towards softer matrix. Arrowhead indicates the neurite with Kinesin-1 accumulation. Scale bar, 20 µm.

## SUPPLEMENTARY MOVIES

**Supplementary Movie 1**. Transient accumulation of Kinesin-1 in neurites of stage 2 neuron. Scale bar 10 µm. This video complements Fig. 2.

**Supplementary Movie 2**. Transient accumulation of Kinesin-1 in neurites stops upon Rac inhibition in stage 2 neuron. Scale bar 10 µm. This video complements Fig. 2.

**Supplementary Movie 3**. Photoactivation of Rac1 directs the selective transport of Kinesin-1 in stage 2 neuron. Scale bar 10 µm. This video complements Fig. 2.

**Supplementary Movie 4**. Selective transport of Kinesin-1 is associated with TRIM46 accumulation in the neurites. Scale bar 10 µm. This video complements Fig. 3.

**Supplementary Movie 5**. Anterograde and retrograde movement of TRIM46 patches in stage 2 neurons. Scale bar 5 µm. This video complements Fig. 3.

**Supplementary Movie 6**. Microtubule network mobility at different stages of neuron development. This video complements Fig. 3.

**Supplementary Movie 7**. Changes of TRIM46 accumulation in neurites reduces during neuron development. This video complements Fig. S7.

**Supplementary Movie 8**. Differential microtubule network mobility is associated with selective Kinesin-1 transport. This video complements Fig. 3.

**Supplementary Movie 9. (**Top)Appearance of microtubules near the plasma membrane upon Rapalog induced microtubule anchoring in U2OS cells. (Bottom) Microtubule movement is reduced after Rapalog induced anchoring to the plasma membrane. This video complements Fig. S8.

**Supplementary Movie 10**. Light induced local microtubule anchoring directs the selective transport of Kinesin-1 in stage 2 neuron. This video complements Fig. 4.

**Supplementary Movie 11**. TRIM46 enrichment in the neurite upon light-induced microtubule anchoring. This video complements Fig. S9.

**Supplementary Movie 12**. Control experiment for light induced microtubule anchoring-Kinesin-1 is not directed to the blue light illuminated neurite. This video complements Fig. S10.

